# The First Implication of Image Processing Techniques on Influenza A Virus Sub-Typing Based on HA/NA Protein Sequences, using Convolutional Deep Neural Network

**DOI:** 10.1101/448159

**Authors:** Reza Ahsan, Mansour Ebrahimi

## Abstract

Increase in influenza A virus host range throughout its evolution has given rise to major concerns worldwide. Although the increasing host range mechanism of the virus is largely unknown; persistent genetic mutations have been blamed as a key factor in the re-organization of the host response and the host range. To uncover the underlying core bases of the two important antigenic proteins of influenza virus (HA and NA), functional data mining and image processing analysis of over 8000 protein sequences of different HA and NA subtypes were performed. Each amino acid sequence in HA or NA proteins sat as a feature or variable and two polynomial datasets were created and subjected into conventional prediction models. The average accuracies of these predictive models for HA subtype classifications ranged from 38.9% for SVM to 87.2% for Decision Tree models. NA subtype classification with conventional prediction models varied from 81.3% to 99.87% for SVM and KNN models, respectively. Then amino acid sequence datasets were converted to binary images; subtypes feature sat as target variable, and target label determined by image processing convolution neural network. The performances of Image processing models (convolutional neural network) on binary images for both HA and NA datasets reached to 100%; and the application of Gabor2 filter decreased the time for the predicting model to reach the best performance for HA subtype; while it increased the epochs time for NA subtype classification.

For the first time ever, converting influenza virus’ HA and NA amino acid sequences into the binary image datasets and their classifications by convolution neural network increased the prediction accuracies and performances to the highest possible point. The finding of this paper paves new avenues for virus classification based on antigenic HA and NA amino acid profiles, and easily classifying and predicting the possible future emerging strains of pandemic influenza.

## Introduction

The ability of influenza virus to increase its host range is a major concern worldwide; resulting in human infection with high mortality rate and widespread pandemic fear with higher morbidity and mortality rates [1]. The last influenza outbreak with a novel avian origin influenza A (H7N9), caused up to 400 000 deaths globally in the first year, and has increased concerns over its pandemic potentials in near future [2]. The emergence of new broad host range of influenza strain with lack of previous host immunity and human to human transmission ability may result in the real pandemic outbreak with millions of fatalities. High frequency of genetic reassortment and antigenic drift, availability of hosting environments and circulating different subtypes of influenza virus for genetic alteration provide the virus suitable setting to generate new highly infectious strains [3].

Based on two surface glycoproteins, haemagglutinin (HA) and neuraminidase (NA), 16 HA subtypes and 9 NA subtypes of influenza virus have been identified. Less than thirty percent of HA and NA amino acids are conserved among all virus subtypes and HA and NA segments are extremely variable in genetic sequences [4]. Its surface HA proteins are the key part in the specificity of influenza virus infection, while during viral releases from host cells, the cleavage of linkage between terminal sialic acid and adjacent galactose is done by NA. Influenza virus A strategy to increase its host range goes through alteration of viral surface proteins. it has been shown that a few amino acid substitutions enabled the virus to transmit via respiratory droplet between ferret. Or single amino acid substitution converted nonlethal strain of influenza to a lethal virus in human; showing the importance of amino acid profiling of surface influenza proteins to monitor the host specificity [5].

Development of algorithms that allow computers to extract the patterns among the data variables is a subfield of artificial intelligence. Machine learning goes through a process of inference and fitting the best model or learning algorithm from examples. The approaches have been widely used in many applications (i.e. pattern recognition, stock market prediction, text and language processing and development of search engines) [6–10]. Multiple sequence alignment, protein structure prediction, gene expression analysis, gene ontology prediction, and molecular classification are other areas of prediction models. Machine learning techniques have achieved great success in biological classifications and evolutionary pattern recognition, including influenza virus host and subtype identification [11, 12].

Deep learning is a branch of machine learning discussion and a set of algorithms that try to model high-level abstract concepts using learning at different levels and layers [13, 14]. The depth, the number of node layers process data to recognize the pattern, is the most distinguished feature of deep learning from conventional neural networks. Training nodes on a distinct set of features based on the output of the previous are the most important characteristic of deep neural networks [15, 16].

Historically, image processing and face recognition tasks have been done successfully by deep neural networks algorithms [17]. These models can be trained to detect objects in pictures far more better than human do [18]. The most common used deep learning network architecture for image analysis is the convolutional neural network (CNN). Pattern matching (convolution) and aggregation (pooling) operations are the basic cores of CNN. Scanning the image by a given pattern and calculating of a match for every position is done at the pixel level. The presence of the pattern in a region determines by pooling (max-pooling), and the region information aggregates into a single number [19, 20]. Most network architectures used in image processing and image analysis have been done by convolution and pooling operations [21].

In this research, for the first time, two important influenza virus A subtypes’ protein sequences (HA and NA) converted into binary images and their subtypes were predicted by developing, training and validating image processing convolution neural network algorithms; and finally their performances compared with the conventional predictive models.

## Material and Methods

The following steps were undertaken as shown in Figure 1.:

i. constructing polynomial datasets of HA & NA amino acids’ sequences;
ii. constructing a binary image datasets of HA & NA amino acids’ sequences;
iii. training and testing conventional predictive models on polynomial datasets;
iv. developing, training and validating the convolution neural network (CNN) to predict the virus subtypes based on images of protein sequences
v. comparing the predictive performances of CNN with conventional predictive models

**Figure 1.**
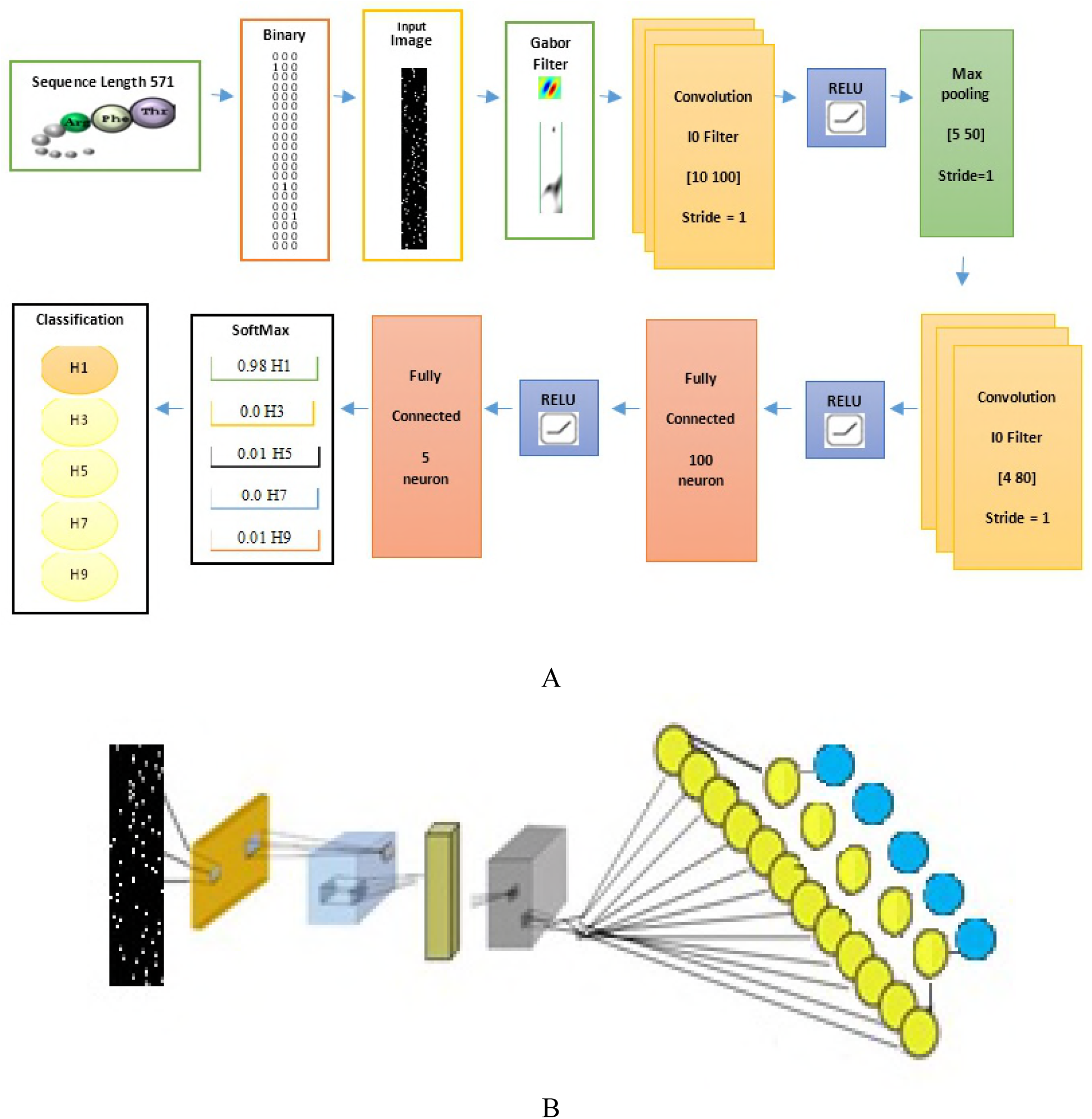
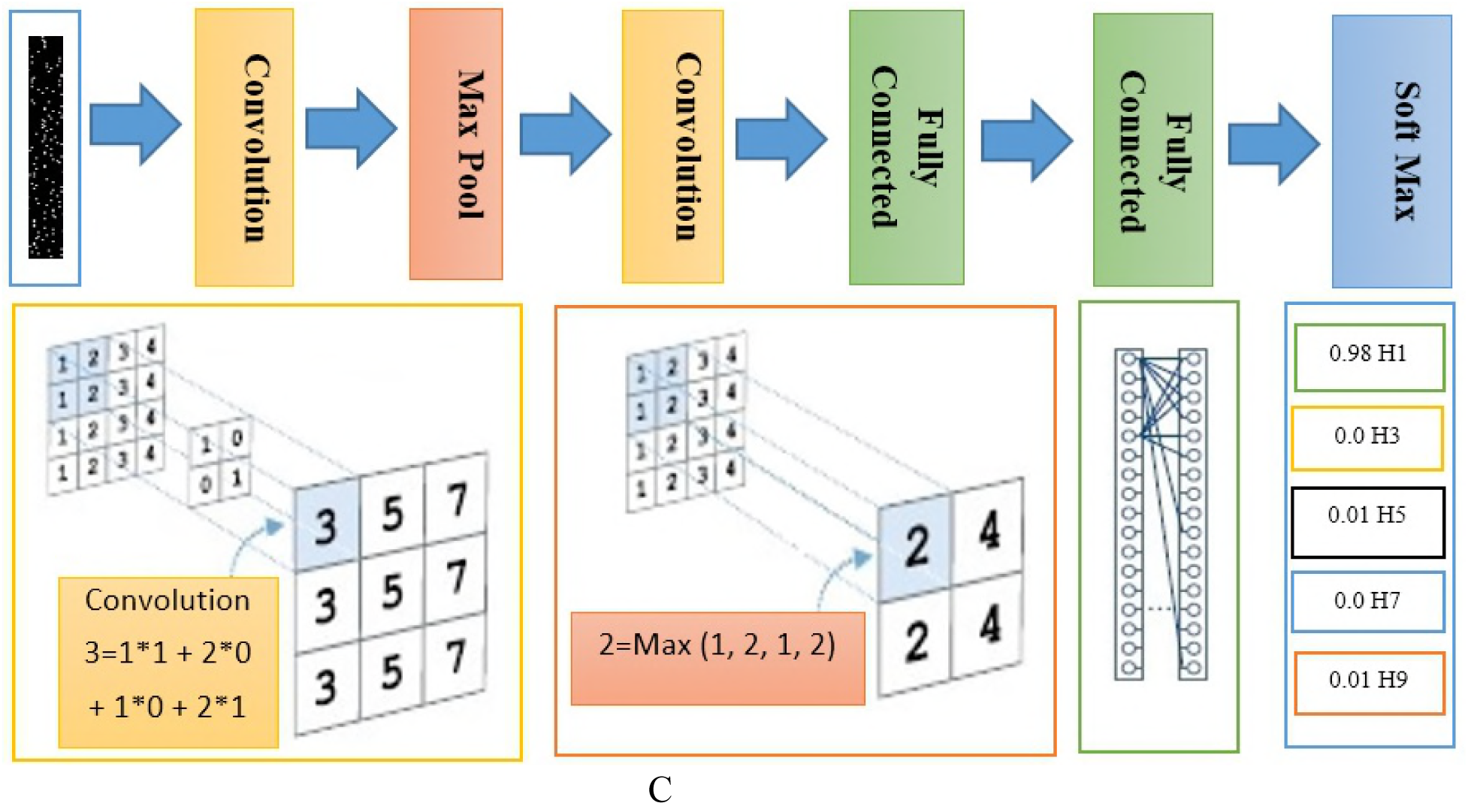
Overview of CNN network topology (A) and various steps taken to convert protein sequences into a binary image and applying Gabor filter to find the right subtype classes (B); the arrangement of the layers in the CNN has been presented in part (C).

### Data architecture and datasets

Sequences of five hemagglutinin (HA) proteins (H1, H3, H4, H5, H9) and four neuraminidase (NA) proteins (N1, N2, N6, and N8) with at least 500 samples in each group extracted from UniProt Protein database (https://www.uniprot.org). The total number of protein sequences for each influenza virus A subtype was 4000 examples. The following datasets generated for each virus subtype:

1. **HA Polynomial Dataset (HAPD)**: Each amino acid position converted to one feature or variable; as the longest HA protein sequence made of 576 amino acids, therefore, 576 features (or column) for each sample created. This dataset contained a matrix of 4000 protein samples and 576 amino acid position as variables.
2. **NA Polynomial Dataset (NAPD)**: The longest NA protein sequence made of 475 amino acids, a polynomial dataset of 4000 rows of NA protein sequences and 475 column of each amino acid variables or features created.
3. **HA Binary Image Dataset (HABID): Regarding the total numbers of amino acids of** 20, to each amino acid letter a digit from 1 to 20 assigned (See Table 1.). The assignment based on *SeqInt = aa2int(SeqChar)* function which converts sequence character of single-letter codes of an amino acid to an integer; based on Table 1 values. Then numeric data of HA sequences converted to the binary image; composed of nineteen 0 and 1. For example, we assigned number 2 to amino acid Arginine (R); therefore its binary numbers would be 01000000000000000000. The final created image made of 20 * 576 binary matrix.
4. **NA Binary Image Dataset (NABID)**: Again, to each amino acid sequence of NA protein a digit between 1 and 20 assigned (as explained above). Each digit converted to a binary image; 1 assigned at the position with a number and for the rest of 19 more spaces, nineteen 0 assigned. The final image dataset made of 20 * 475 binary matrix.

**Table 1.** Conversion table used for assigning a digit to each amino acid letter.

| **Amino Acid** | **Code** | **Integer** |
| --- | --- | --- |
| Alanine | A | 1 |
| Arginine | R | 2 |
| Asparagine | N | 3 |
| Aspartic acid<br>(Aspartate) | D | 4 |
| Cysteine | C | 5 |
| Glutamine | Q | 6 |
| Glutamic acid | E | 7 |
| Glycine | G | 8 |
| Histidine | H | 9 |
| Isoleucine | I | 10 |
| Leucine | L | 11 |
| Lysine | K | 12 |
| Methionine | M | 13 |
| Phenylalanine | F | 14 |
| Proline | P | 15 |
| Serine | S | 16 |
| Threonine | T | 17 |
| Tryptophan | W | 18 |
| Tyrosine | Y | 19 |
| Valine | V | 20 |
| Unknown character | ? | 0 |

### Conventional Machine Learning Predictive Models

The polynomial datasets (HAPD and NAPD) imported into MATLAB (*R2018b, 9.4.0.813654, MathWorks, 1 Apple Hill Drive Natick, MA, USA);* the type of HA (H1, H3, H4, H5 and H9 classes) or NA (N1, N2, N6 and N8 classes) sat as target or label variable. The following conventional classification learner models applied on both datasets: Decision Tree (Fine Tree, Medium Tree, Coarse Tree), Discriminant Linear analysis, Support Vector Machine (SVM: Linear, Quadratic, Cubic, Cubic, Fine Gaussian, Medium Gaussian, Coarse Gaussian), Nearest Neighborhood (KNN: Fine KNN, Medium KNN, Coarse KNN, Weighted KNN) and Ensemble (Boosted Tree, Bagged Trees, Subspace KNN, RUSBoosted Trees) classifiers.

To make the results comparable, no cross-validation approach selected; data divided into 90% and 10% parts and each model trained on nine parts and validated (tested) on the tenth part. The performance or the accuracy of each model in predicting the right class of HA or NA subtypes computed.

### Image Processing Convolution Neural Network (CNN)

A CNN made of three major layers: convolution, pooling, and fully connected layers; each layer does a special task. There are two stages for training in CNN; feed-forward and back-propagation. In the first stage, the image fed into the network; This action is nothing but a multiplication of the point between the input image matrix and the filter matrix in each convolution layer. The layer searches for high-level features extracted from raw data; looking for meaningful objects and; no decision is being made at this stage. Flatting these features at the end of the network and connecting them to two fully-connected layers is usually a cheap computational load method for learning the nonlinear components of these features. The dimensions of the weight matrix to produce the number of neurons required in the all-connected layer are equal to the product of multiplying the number of these neurons in the number of neurons in their previous layer. The RELU conversion function is used to zero the negative values of the resulting matrix. The integration layer is the layers that are placed after the convolutional layer and merely reduce the size of the data. There are various mechanisms for it, most notably Max Pooling. In this mechanism, the windows on the resulting matrix are applied to the previous step and moves with a certain step. Its task is to place the maximal numbers in the window instead of the numbers. The Softmax layer takes the output of the previous layer and converts it into the probability distribution of the classes. The output of the Softmax function is a number between 0 and 1, and the sum of these numbers is one. Then the network output is calculated. Here, for the purpose of setting network parameters, i.e., the values of convolutional layer filters and weight matrices of fully connected layers, or in other words, the network training, the output is used to calculate the network error rate. To do this, the network compared output using an error function with the correct response, and the error rate is calculated. The next step is based on the calculated error rate of the backpropagation stage. In this step, the gradient of each parameter is calculated according to the law of the chain rule, and all parameters are changed according to the effect on the error generated in the network. After the parameters are updated, the next phase of the feed-forward starts. After completing the correct number of these steps, the network training ends.

Two binary image datasets of HA and NA imported into MATLAB (*R2018b, 9.4.0.813654, MathWorks, 1 Apple Hill Drive Natick, MA, USA*); 90% of data in each dataset sat as training set and the last 10% as testing or validation set. To find the best algorithm, various convolution neural networks’ architectures were examined. To extract the meaningful features, the first layer filter sat as 10 × 100. RELU function used to remove negative figures. To reduce image dimensions, 5 × 50 windowing filter applied just once. The second convolution layer sat as a smaller layer of 4 × 80. The first layer of the network had 100 neurons while the second fully connected layer made of just 5 neurons; based on 5 or 4 classes of HA or NA influenza viruses’ subtypes, respectively. For probability distribution and network output class presentation, Softmax and classification layers were used. Figure 1 shows the convolution network architecture and layers’ architecture.

To enhance the classification accuracy for each class; GABOR2 filter is applied with 90-degree angles on horizontal images at the preprocessing stage on the binary image.

## Results

### Prediction Model

Conventional machine based predictive models were trained and validated on polynomial protein sequences of influenza virus A subtypes (HA and NA) and compared with the same dataset when it converted into a binary image dataset and analyzed by the image processing predictive model (convolutional neural network - CNN). No cross-validation method selected for CNN, and to make the results comparable, no cross-validation methods were also chosen for the conventional predictive models (Decision Tree - Fine Tree, Medium Tree, Coarse Tree), Discriminant Linear analysis, Support Vector Machine (SVM: Linear, Quadratic, Cubic, Fine Gaussian, Medium Gaussian, Coarse Gaussian), Nearest Neighborhood (KNN: Fine KNN, Medium KNN, Coarse KNN, Weighted KNN) and Ensemble (Boosted Tree, Bagged Trees, Subspace KNN, RUSBoosted Trees classifiers).

### Conventional Predictive Models

1. **HA Polynomial Dataset (HAPD)**: The best average accuracy in predicting the right class of HA subtype gained with Decision Tree (87%) prediction model; followed by Ensemble and KNN models. The lowest accuracy gained by SVM prediction models..
2. **NA Polynomial Dataset (NAPD)**: The lowest average performance gained by SVM (81.3%), while the performance of Ensemble models were around 90%. The best prediction of 99.8% gained by KNN models.
3. **HA Binary Image Dataset (HABID)**: As seen in Figure 2, the performances of CNN model in predicting the five classes of HA proteins based on their binary image dataset were outstanding (100%). All 10% validation test samples were exactly predicted in their right class and nothing left out. When Gabor2 filter added to the prediction procedure, the performance did not change but the model reached the highest possible accuracy in a shorter time (epoch 5 compared to epoch 12) (Figure 3).
4. **NA Binary Image Dataset (NABID)**: The performance of image processing model in classifying NA protein classes of influenza virus A again reached the top possible figure of 100% in just 3 epochs. When Gabor2 filter applied, the model reached to the top accuracy in 7 epochs (Figure 4.)

**Figure 2.**
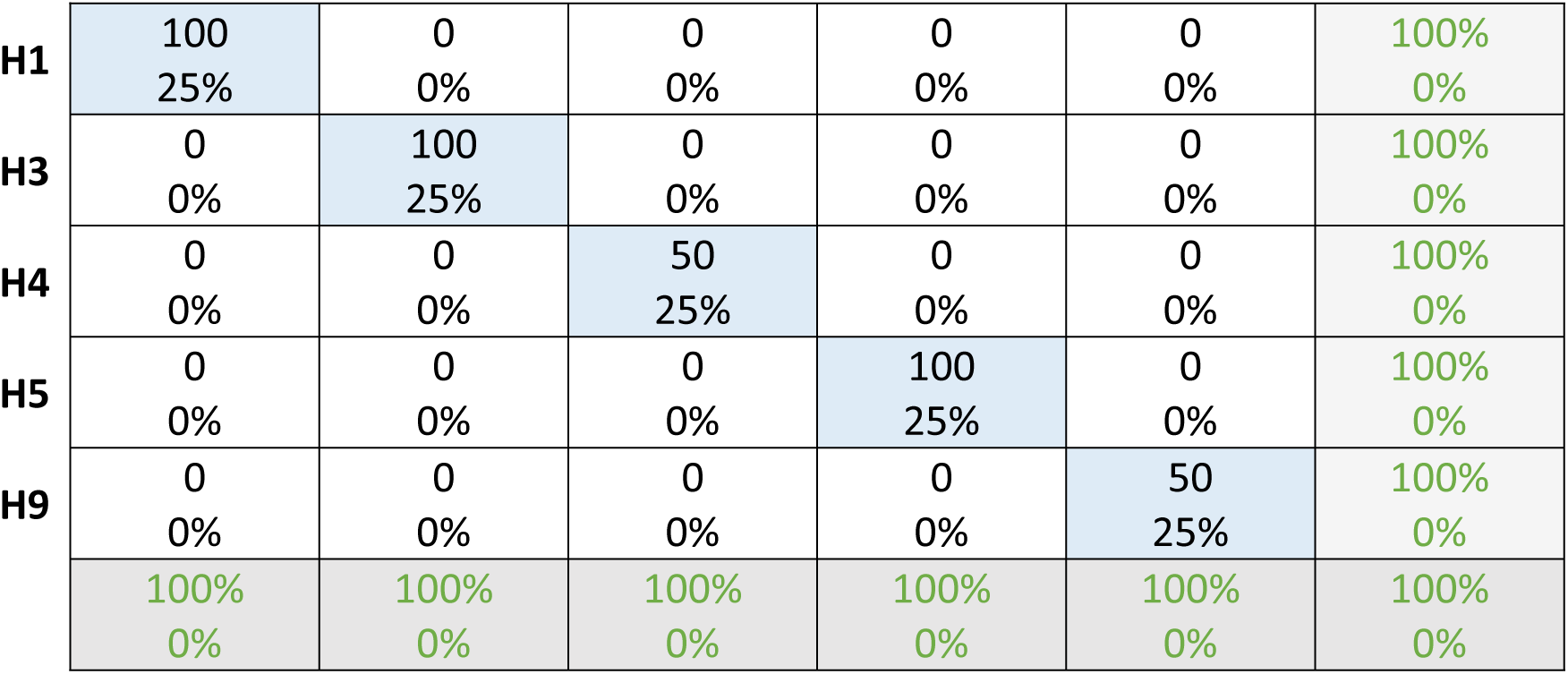
Confusion Matrix of CNN predictive models ran on binary image dataset of HA subtype protein sequences without Gabor2 filter.

**Figure 3.**
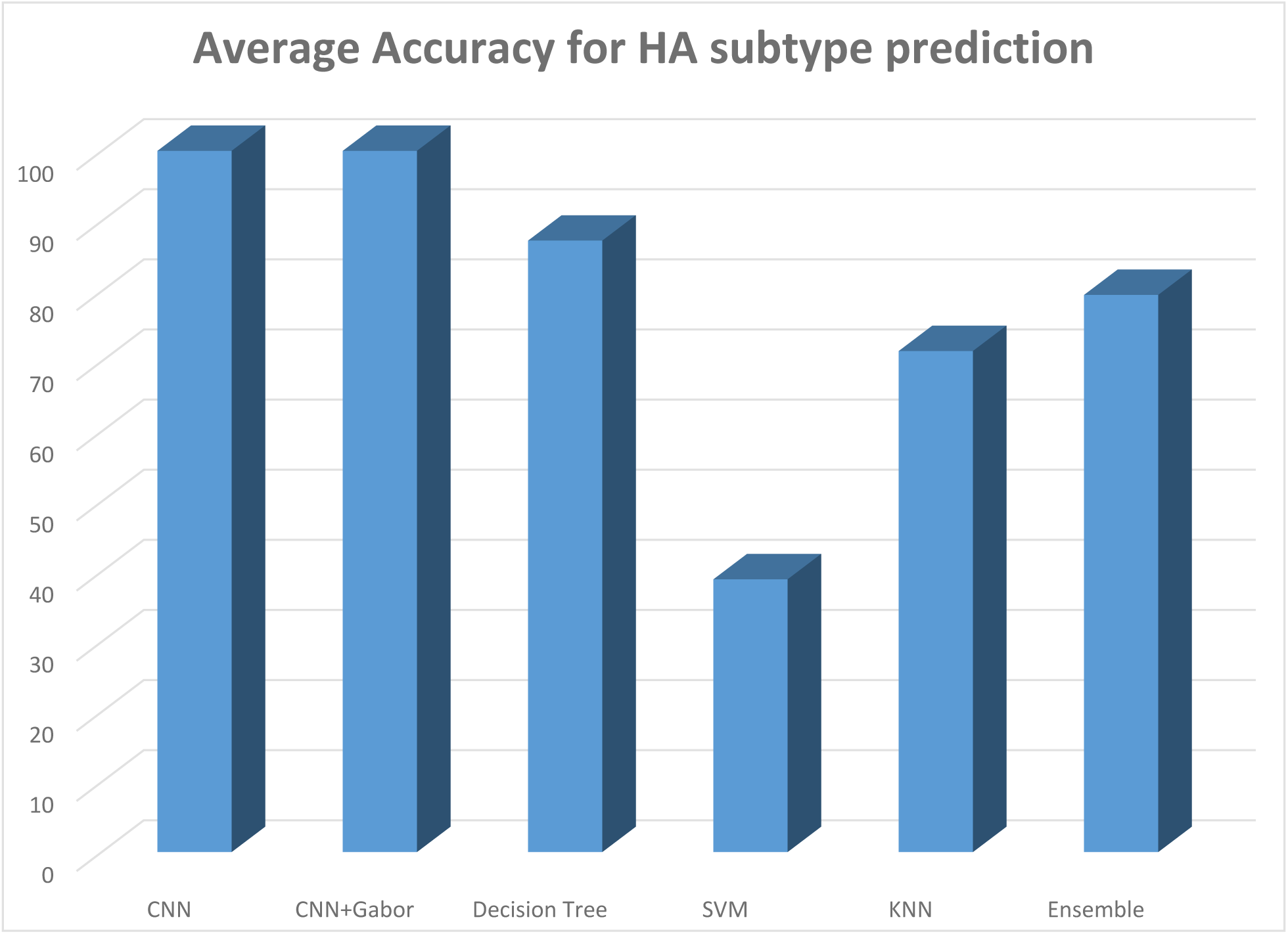
The percentage of accuracies of various prediction models (conventional and CNN) in predicting the right class of HA (H1 - H4).

**Figure 4.**
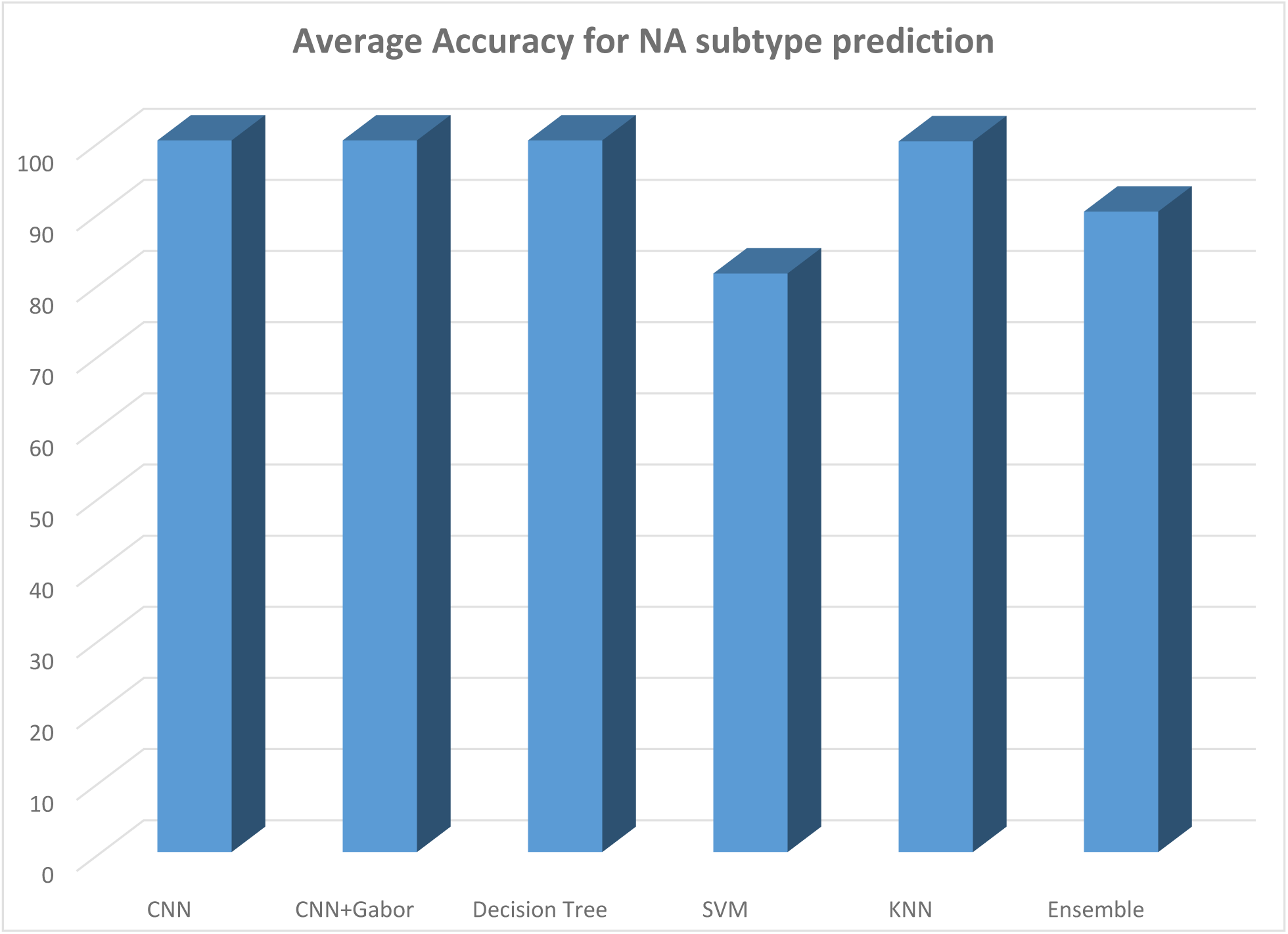
The percentage of accuracies of various prediction models (conventional and CNN) in predicting the right class of NA (N1 - N4).

## Discussion

Influenza A pandemics of the last century (H1N1, H2N2, and H3N2), avian H5N1 and recent H7N9 have been infecting the human with high mortality rates [22]. Multiple gene reassortant of viruses from various sources (such as swine, avian and even human populations) have been blamed for the H1N1 pandemic in 2009 and novel reassortant of H7N9 in 2013; expecting more pandemics to happen in near future due to high reassortment capacity of the virus [23]. Therefore understanding the biological bases of influenza virus A evolution and subtype differentiation is critical. The virus undergoes rapid evolution in the antigenic regions when subjected to host immune response and crosses the host species barrier. Altering rapid short amino acid patterns enable the virus to avoid time-consuming major changes in proteins, and so far, the strategy of rapid structural amino acid alteration has worked well for the virus [24, 25].

In line with our previous results [12, 26], to find the most important features which can be used to distinguish between the virus subtypes, attribute weighting models applied on the polynomial dataset. The results showed that, for HA subtypes, two positions (251 and 541) were the most important positions weighed the highest possible weight higher than 0.9 by nine models. Amino acids’ positions of 185, 248, 317 and 447 were the most important features in distinguishing between four classes of influenza virus A subtypes based on NA protein sequences. As each attribute weighting model uses a specific statistical pattern to find the most important features, when any features appointed by ninety percent of the models as important, it means that those features are highly distinctive among virus subtypes.

As we previously showed, influenza virus can acquire additional host specificity of another subtype just by a small change in amino acid sequence; increasing the chance of host range and survival by less energy consumption and minimum change in protein sequences [22].

Recently, there has been a lot of interest in image processing techniques and they have been widely used in many different fields. Face recognition and license plate detection are two of the widest areas of research in the application of CNN [27, 28]. Medical image pattern recognition is another attractive field of research for image processing and machine-based prediction neural networks [29]. CNN architecture and image processing with texture feature parameters have been used in the classification of masses and normal tissue on mammograms with 90% accuracy [30, 31]. Detection of lung nodules, classification of radiograms and bone fractures has attracted many research attractions [32, 33].

This is the first successful attempt in modeling and prediction of influenza virus A subtypes based on image processing algorithms. Highly accurate models developed, trained and validated by analyzing images made of amino acid sequences of antigenic proteins of the influenza virus. The models were able to convert the raw protein sequences into images in a few short seconds and then apply the image processing techniques to identify the prominent key differences between virus subtypes’ images.

The performances or accuracies of conventional predictive models (as we named in this paper) have been compared with newly developed image processing CNN algorithms of protein sequences. For the first time, we created an image based on amino acids sequences, the binary image then fed into the CNN algorithms to compute the relationship between the image and the influenza A virus subtype. Instead of creating polynomial datasets, quickly each sequence converted into an image and the CNN model employed to process the images, learn the relationships, then tested and validated. The outcome was fascinating, a machine learning tool based on image processing capabilities whit nearly hundred percent precision in classifying influenza A virus subtypes.

The performances of conventional predictive models varied, from 35% to 99% The findings are in line with previously published works. Although we were able to reach 99% accuracy with Naïve Bayes model in predicting the HA subtype [22], that dataset created based on thousands of physic-o-chemical features of proteins, not protein sequence. But here, raw amino acid sequences fed into the model; now, the developed algorithm can be directly used to predict the type of HA or NA proteins with the highest possible accuracy.

Although the number of amino acid sequences in HA or NA subtype classes in the prepared datasets was not equal (the number of some HA or NA classes were twice the others), the model did a great job in predicting the right virus subtype class. Therefore, the developed algorithm can be used to predict influenza virus A subtypes with any number of protein sequences; making the model suitable for asymmetric datasets.

This research approach will serve as a fundamental base for future studies on clustering influenza virus A based on raw protein or genetic sequences of other virus segments (such as M2E), or even the whole virus genome or proteome. Applying this approach to other viruses can facilitate classification based on genomic or proteomic sequences. The novel methods developed in this research can be embedded in software or even web-based applications to predict the type of newly emerged influenza virus based on sequences. Combinations of this research results with future studies on antigen and antibody interaction measurements by ELISA or Western Blot can result in vaccine breakdown and developing new efficient vaccines based on genomic and proteomic sequences. In addition, developed models can be used in future investigations on other influenza virus amino acids compositions (such as M2E) and their possible roles in viral antigenicity. The findings clearly suggest the viral amino acid profiles of the influenza virus as potential features to monitor host specificity [34].

Study of the relationship between protein sequence profiles and disease states plays an important role in clinical and biological applications [35]. A deep learning algorithm can be the most suitable choice of predicting the disease status based on genomic or proteomic data. This method extracts the feature in a hierarchy of layers through nonlinear functions. The input of each layer is the output of the previous layer, and its training can be either observer or non-observer. In fact, the single layer of the hidden layer in the neural network is replaced with a large number (deep) of the layer. Convulsion neural networks are one of the most important learning methods in which several layers are trained in a powerful way. This method is very efficient and is one of the most commonly used methods in various computer vision applications [36].

## Conclusion

Computer-based predictive models have opened new vistas in medical analyses and diagnostic tests and their implications in these fields are growing rapidly. In this research, we developed and applied image recognition convolution deep learning neural network algorithms to distinguish between five different HA and four NA subtypes of influenza virus A.

For the first time, we converted and transformed protein sequences into images and by optimizing the image processing filters, and we were able to classify the virus subtypes with 100% accuracies. Comparing the results of the developed method with the conventional predictive models showed this approach was more efficient, accurate and less time-consuming.

As this method can be used to quickly convert and compare whole genome and proteome sequences of healthy and unhealthy people into images, it really opens new analytical approaches. The method can be used easily to compare any genome, transcriptome or proteome of organisms at the different situations,

## References

1. Bucher J, Boelle PY, Hubert D, Lebourgeois M, Stremler N, Durieu I, et al. Lessons from a French collaborative case-control study in cystic fibrosis patients during the 2009 A/H1N1 influenza pandemy. BMC Infect Dis. 2016;16:55. doi: 10.1186/s12879-016.2-1352-PubMed PMID: 26830335; PubMed Central PMCID: PMCPMC4736161.

2. Kim SM, Kim YI, Pascua PN, Choi YK. Avian Influenza A Viruses: Evolution and Zoonotic Infection. Semin Respir Crit Care Med. 2016;37(4): 501–11. doi: 10.1055/s-0036-1584953. PubMed PMID:27486732:

3. Cueno ME, Suzuki I, Shimotomai S, Yokoyama T, Nagahisa K, Imai K. Structural comparison among the 2013-2017 avian influenza A H5N6 hemagglutinin proteins: A computational study with epidemiological implications. J Mol Graph Model. 2018;79:1.91–85doi: 10.1016/j.jmgm.2017.11.013. PubMed PMID: 29220671.

4. Houghton R, Ellis J, Galiano M, Clark TW, Wyllie S. Haemagglutinin and neuraminidase sequencing delineate nosocomial influenza outbreaks with accuracy equivalent to whole genome sequencing. J Infect. 2017;74(4): 377–84. doi: 10.1016/j.jinf.2016.12.015. PubMed PMID: 28104386.

5. Bera BC, Virmani N, Kumar N, Anand T, Pavulraj S, Rash A, et al. Genetic and codon usage bias analyses of polymerase genes of equine influenza virus and its relation to evolution. BMC Genomics. 2017;18(1): 652. doi: 10.1186/s12864-017-4063-1. PubMed PMID: 28830350; PubMed Central PMCID: PMCPMC5568313.

6. Pashaiasl M, Khodadadi K, Kayvanjoo AH, Pashaei-Asl R, Ebrahimie E, Ebrahimi M. Unravelling evolution of Nanog, the key transcription factor involved in self-renewal of undifferentiated embryonic stem cells, by pattern recognition in nucleotide and tandem repeats characteristics. Gene. 2016;578(2): 194–204. doi: 10.1016/j.gene.2015.12.023. PubMed PMID: 26687709.

7. Banik S, Khodadad Khan AF, Anwer M. Hybrid machine learning technique for forecasting Dhaka stock market timing decisions. Comput Intell Neurosci. 2014;2014:318524. doi: 10.1155/2014/318524. PubMed PMID: 24701205; PubMed Central PMCID: PMCPMC3950395.

8. Pyo S, Lee J, Cha M, Jang H. Predictability of machine learning techniques to forecast the trends of market index prices: Hypothesis testing for the Korean stock markets. PLoS One. 2017;12(11):e0188107. doi: 10.1371/journal.pone.0188107. PubMed PMID: 29136004;PubMed Central PMCID: PMCPMC5685607.

9. Badal VD, Kundrotas PJ, Vakser IA. Natural language processing in text mining for structural modeling of protein complexes. BMC Bioinformatics. 2018;19(1): 84. doi: 10.1186/s12859-018-2079-4. PubMed PMID: 29506465;PubMed Central PMCID: PMCPMC5838950.

10. Voulodimos A, Doulamis N, Bebis G, Stathaki T. Recent Developments in Deep Learning for Engineering Applications. Comput Intell Neurosci. 2018;2018:8141259. doi: 10.1155/2018/8141259. PubMed PMID: 29861713; PubMed Central PMCID: PMCPMC5971264.

11. Bakhtiarizadeh MR, Moradi-Shahrbabak M, Ebrahimi M, Ebrahimie E. Neural network and SVM classifiers accurately predict lipid binding proteins, irrespective of sequence homology. J Theor Biol. 2014;356:213–22. doi: 10.1016/j.jtbi.2014.04.040. PubMed PMID: 24819464.

12. Ebrahimi M, Aghagolzadeh P, Shamabadi N, Tahmasebi A, Alsharifi M, Adelson DL, et al. Understanding the undelaying mechanism of HA-subtyping in the level of physic-chemical characteristics of protein. PLoS One. 2014;9(5):e96984. doi: 10.1371/journal.pone.0096984. PubMed PMID: 24809455; PubMed Central PMCID: PMCPMC4014573.

13. Biswas M, Kuppili V, Araki T, Edla DR, Godia EC, Saba L, et al. Deep learning strategy for accurate carotid intima-media thickness measurement: An ultrasound study on Japanese diabetic cohort. Comput Biol Med. 2018;98:100–17. doi: 10.1016/j.compbiomed.2018.05.014. PubMed PMID: 29778925.

14. Cao C, Liu F, Tan H, Song D, Shu W, Li W, et al. Deep Learning and Its Applications in Biomedicine. Genomics Proteomics Bioinformatics. 2018;16(1): 17–32. doi: 29522900; PubMed Central PMCID: PMCPMC6000200.

15. Jing Y, Bian Y, Hu Z, Wang L, Xie XS. Deep Learning for Drug Design: an Artificial Intelligence Paradigm for Drug Discovery in the Big Data Era. AAPS J. 2018;20(3): 58. doi: 10.1208/s12248-018-0210-0. PubMed PMID: 29603063.

16. Lan K, Wang DT, Fong S, Liu LS, Wong KKL, Dey N. A Survey of Data Mining and Deep Learning in Bioinformatics. J Med Syst. 2018;42.139:(8) doi: 10.1007/s10916-018-1003-9. PubMed PMID: 29956014.

17. Rodellar J, Alferez S, Acevedo A, Molina A, Merino A. Image processing and machine learning in the morphological analysis of blood cells. Int J Lab Hematol. 2018;40 Suppl 1: 46–53. doi: 10/.1111ijlh.12818. PubMed PMID: 29741258.

18. Yang L, Wong CM, Chiu SS, Cowling BJ, Peiris JS. Estimation of excess mortality and hospitalisation associated with the 2009 pandemic influenza. Hong Kong Med J. 2018;24 Suppl 6(5): 19–22. PubMed PMID: 30229731.

19. Mall S, Brennan PC, Mello-Thoms C. Modeling visual search behavior of breast radiologists using a deep convolution neural network. J Med Imaging (Bellingham). 2018;5(3): 035502. doi: 10.1117/1.JMI.5.3.035502. PubMed PMID: 30128329; PubMed Central PMCID:PMCPMC6086967.

20. Bell JS, Wilson RI. Behavior Reveals Selective Summation and Max Pooling among Olfactory Processing Channels. Neuron. 2016;91(2): 425–38. doi: 10.1016/j.neuron.2016.06.011. PubMed PMID: 27373835; PubMed Central PMCID: PMCPMC5217404.

21. Angermueller C, Parnamaa T, Parts L, Stegle O. Deep learning for computational biology. Mol Syst Biol. 2016;12(7): 878. doi: 10.15252/msb.20156651. PubMed PMID: 27474269; PubMed Central PMCID: PMCPMC4965871.

22. Hemmatzadeh F, Keyvanfar H, Hasan NH, Niap F, Bani Hassan E, Hematzade A, et al. Interaction between Bovine leukemia virus (BLV) infection and age on telomerase misregulation. Vet Res Commun. 2015;39(2): 97–103. doi: 10.1007/s11259–015–9629–2. PubMed PMID: 25665900.

23. Kargarfard F, Sami A, Mohammadi-Dehcheshmeh M, Ebrahimie E. Novel approach for identification of influenza virus host range and zoonotic transmissible sequences by determination of host-related associative positions in viral genome segments. BMC Genomics. 2016;17(1): 925. doi: 10.11/86s12864-016-3250-9. PubMed PMID: 27852224; PubMed Central PMCID: PMCPMC5112743.

24. Bangaru S, Nieusma T, Kose N, Thornburg NJ, Finn JA, Kaplan BS, et al. Recognition of influenza H3N2 variant virus by human neutralizing antibodies. JCI Insight. 2016;1(10) doi: 10.1172/jci.insight.86673. PubMed PMID: 27482543; PubMed Central PMCID: PMCPMC4962875.

25. Clark AM, Nogales A, Martinez-Sobrido L, Topham DJ, DeDiego ML. Functional Evolution of Influenza Virus NS1 Protein in Currently Circulating Human 2009 Pandemic H1N1 Viruses. J Virol. 2017;91(17). doi: 10.1128/JVI.00721-17. PubMed PMID: 28637754; PubMed Central PMCID: PMCPMC5553169.

26. Kargarfard F, Sami A, Ebrahimie E. Knowledge discovery and sequence-based prediction of pandemic influenza using an integrated classification and association rule mining (CBA) algorithm. J Biomed Inform. 2015;57:181–8. doi: 10.1016/j.jbi.2015.07.018. PubMed PMID: 26232668.

27. Phillips PJ, Yates AN, Hu Y, Hahn CA, Noyes E, Jackson K, et al. Face recognition accuracy of forensic examiners, superrecognizers, and face recognition algorithms. Proc Natl Acad Sci U S A. 2018;115(24): 6171–6. doi: 10.1073/pnas.1721355115. PubMed PMID: 29844174; PubMed Central PMCID: PMCPMC6004481.

28. Nguyen DT, Pham TD, Baek NR, Park KR. Combining Deep and Handcrafted Image Features for Presentation Attack Detection in Face Recognition Systems Using Visible-Light Camera Sensors. Sensors (Basel). 2018;18(3). doi: 10.3390/s18030699. PubMed PMID: 29495417; PubMed Central PMCID: PMCPMC5876704.

29. Bai W, Sinclair M, Tarroni G, Oktay O, Rajchl M, Vaillant G, et al. Automated cardiovascular magnetic resonance image analysis with fully convolutional networks. J Cardiovasc Magn Reson. 2018;20(1): 65. doi: 10.1186/s12968-018-0471-x. PubMed PMID: 30217194; PubMed Central PMCID: PMCPMC6138894.

30. Ribli D, Horvath A, Unger Z, Pollner P, Csabai I. Detecting and classifying lesions in mammograms with Deep Learning. Sci Rep. 2018;8(1): 4165. doi: 10.1038/s41598-018-22437-z. PubMed PMID: 29545529; PubMed Central PMCID: PMCPMC5854668.

31. Ma W, Zhao Y, Ji Y, Guo X, Jian X, Liu P, et al. Breast Cancer Molecular Subtype Prediction by Mammographic Radiomic Features. Acad Radiol. 2018. doi: 10.1016/j.acra.2018.01.023. PubMed PMID: 29526548.

32. Nishio M, Sugiyama O, Yakami M, Ueno S, Kubo T, Kuroda T, et al. Computer-aided diagnosis of lung nodule classification between benign nodule, primary lung cancer, and metastatic lung cancer at different image size using deep convolutional neural network with transfer learning. PLoS One. 2018;13(7):e0200721. doi: 10.1371/journal.pone.0200721. PubMed PMID: 30052644; PubMed Central PMCID: PMCPMC6063408.

33. Ali I, Hart GR, Gunabushanam G, Liang Y, Muhammad W, Nartowt B, et al. Lung Nodule Detection via Deep Reinforcement Learning. Front Oncol. 2018;8:108. doi: 10.3389/fonc.2018.00108. PubMed PMID: 29713615; PubMed Central PMCID: PMCPMC5912002.

34. Schepens B, De Vlieger D, Saelens X. Vaccine options for influenza: thinking small. Curr Opin Immunol. 2018;53:22–9. doi: 10.1016/j.coi..2018.03.024. PubMed PMID: 29631195.

35. Yamada KD, Kinoshita K. De novo profile generation based on sequence context specificity with the long short-term memory network. BMC Bioinformatics. 2018;19(1): 272. doi: 10.1186/s12859-018-2284-1. PubMed PMID: 300;21530PubMed Central PMCID: PMCPMC6052547.

36. Schmidhuber J. Deep learning in neural networks: an overview. Neural Netw. 2015;61:85–117. doi: 10.1016/j.neunet.2014.09.003. PubMed PMID: 25462637.

